# Digital therapeutics for distributed response to global pandemics

**DOI:** 10.1101/444851

**Authors:** Adar Hacohen, Reuven Cohen, Sol Efroni, Baruch Barzel, Ido Bachelet

**Author notes:** Address for correspondence: Department of Mathematics, Bar-Ilan University, Ramat-Gan, 52900, Israel.

## Abstract

Despite advances in the development of drugs and vaccines, the spread of infectious diseases remains an imminent threat to our global health, in extreme cases potentially having detrimental consequences. At present our response to this threat is based on physically distributing therapeutic material, which utilizes the same transportation networks that support the spread of the infectious agent itself. Such competition is at risk of failure in the face of a rapidly spreading pathogen, especially given the inevitable delay from the initial outbreak to the development and execution of our response. Moreover, based on our existing transportation networks, we show that such physical distribution is intrinsically inefficient, leading to an uneven concentration of the therapeutic within a small fraction of destinations, while leaving the majority of the population deprived. This suggests that outrunning a virulent epidemic can only be achieved if we develop a mitigation strategy that bypasses the existing distribution networks of biological and chemical material. Here we propose such a response, utilizing *digitizable* therapeutics, which can be distributed as digital sequence files and synthesized on location, exposing an extremely efficient mitigation scheme that systematically outperforms physical distribution. Our proposed strategy, based for example on nucleic acid therapeutics, is plausibly the only viable mitigation plan, based on current technology, that can face a violently spreading pathogen. Complementing the current paradigm, which ranks drugs based on efficacy, our analysis demonstrates the importance of balancing efficacy with distributability, finding that in some cases the latter plays the dominant role in the overall mitigation efficiency.

---

When studied from the angle of its worst-case scenario, surviving a highly infectious pandemic depends on a competition between the infectious pathogen and the therapeutic technology, each racing to reach the majority of the population first. This competition confronts us with several challenges: (i) the inevitable response time *t*_R_ required for us to instigate a mitigation plan places the pathogen at a potentially significant spreading advantage; (ii) while the pathogen reproduces as it spreads^1–3^, a therapy must be manufactured and shipped from one or few sources, whose production and distribution capacity may be limited^4–12^; (iii) the dissemination of antibiotics or vaccines can be hindered by various external factors, such as geopolitical and socio-economical constraints^13–16^, which have little effect on the propagation of bacteria and viruses. But even if such factors are eliminated, *e.g.*, assuming that an effective cure already exists, stockpiled in sufficient quantities and benefits from worldwide cooperation in its distribution, it would still have to outrun the pathogen, competing along the same routes of dissemination as the epidemic, *i.e.* the international transportation networks^17–23^. For a rapidly progressing epidemic, such competition may fail, having detrimental consequences in terms of human life.

It seems, therefore, that the only viable strategy is to severely intervene in international mobility, quarantining airports, restricting travel and effectively eliminating the routes supporting the viral spread^24–26^, reserving them strictly for the distribution of the therapeutic agent. Such major interventions, however, may result in a significant economic burden and major political stress, indeed – *a lesser of two evils* – but still a potentially hurtful toll on global stability.

Hence, we offer to break this gridlock by focusing on therapies that can spread via alternative, intrinsically more efficient, routes compared to those of physical transport. Such alternative is achievable if the therapeutic agent can be converted into digital information, handled and distributed as data, then locally *printed*, *i.e.* synthesized, at its designated destination (**Box I**). Indeed, the Internet provides infrastructure to distribute data extremely efficiently, outperforming any distribution network that mobilizes physical commodities. Hence, using this strategy, a *digitizable* therapeutic agent will compete within time-scales that are orders of magnitude shorter than those characterizing even the most virulent epidemics, benefiting from the effectively immediate transmission of data vs. the restricted physical medium on which pathogens propagate^27^.

Our analysis further indicates that digital distribution, followed by local synthesis, leads to homogeneous spread, as opposed to centralized distribution, where few destinations benefit from superfluous quantities of the therapeutic, while others remain underserved. Such homogeneous availability not only adheres to social values, but also, in the face of a globally spreading epidemic, leads to optimal utility, where similar production capacities translate to significantly more lives saved. Therefore, even if we disregard the difference in the transport time scales, the fact that digitally distributed therapies are locally synthesized and hence evenly disseminated, provides an intrinsic advantage, that can potentially reduce mortality, in some cases by orders of magnitude.

This calls for a change in the current paradigm of classification and prioritization of therapeutics. Our current approach focuses mainly on the therapeutic efficacy^28,29^, *i.e.* how efficiently the biological/chemical agent cures the disease. We relate little weight, however, to the agent’s chemical classification – *e.g.*, whether it is a small molecule, a protein, or a nucleic acid, as, indeed, these details seem marginal as long as it overcomes the lethal pathogen. However, these distinctions become crucial if we consider *digital-distributability*, as not all molecular media are digitizable. At present, the therapeutic family that optimally satisfies both functionality and distributability is represented by polymer sequences^47^, preferably short, of nucleic acids or peptides, which can be disseminated digitally, as sequence files, and then synthesized on location (**Box I**).

## RESULTS

To demonstrate the potential utility of our proposed digital response we consider a highly contagious pandemic spreading globally through air-travel, under the susceptible-infected-removed (SIR) epidemic model^30–32^ (**Box II**). We used empirical data on human aviation to evaluate the flux of passengers between *N* = 1,292 local populations, each with *M_n_* individuals (*n* = 1, …, *N*), and quantified the death toll of the disease by the global mortality

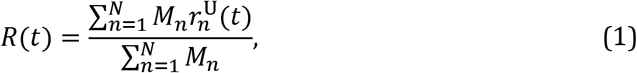

where 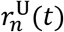 is the fraction of *removed untreated* individuals from the *n*th population (node *n*). For a lethal epidemic, absent any treatment, we have *R* ≡ *R* (*t* → ∞) → 1, representing the demise of the entire population (Fig. 1a, grey).

**Figure 1.**
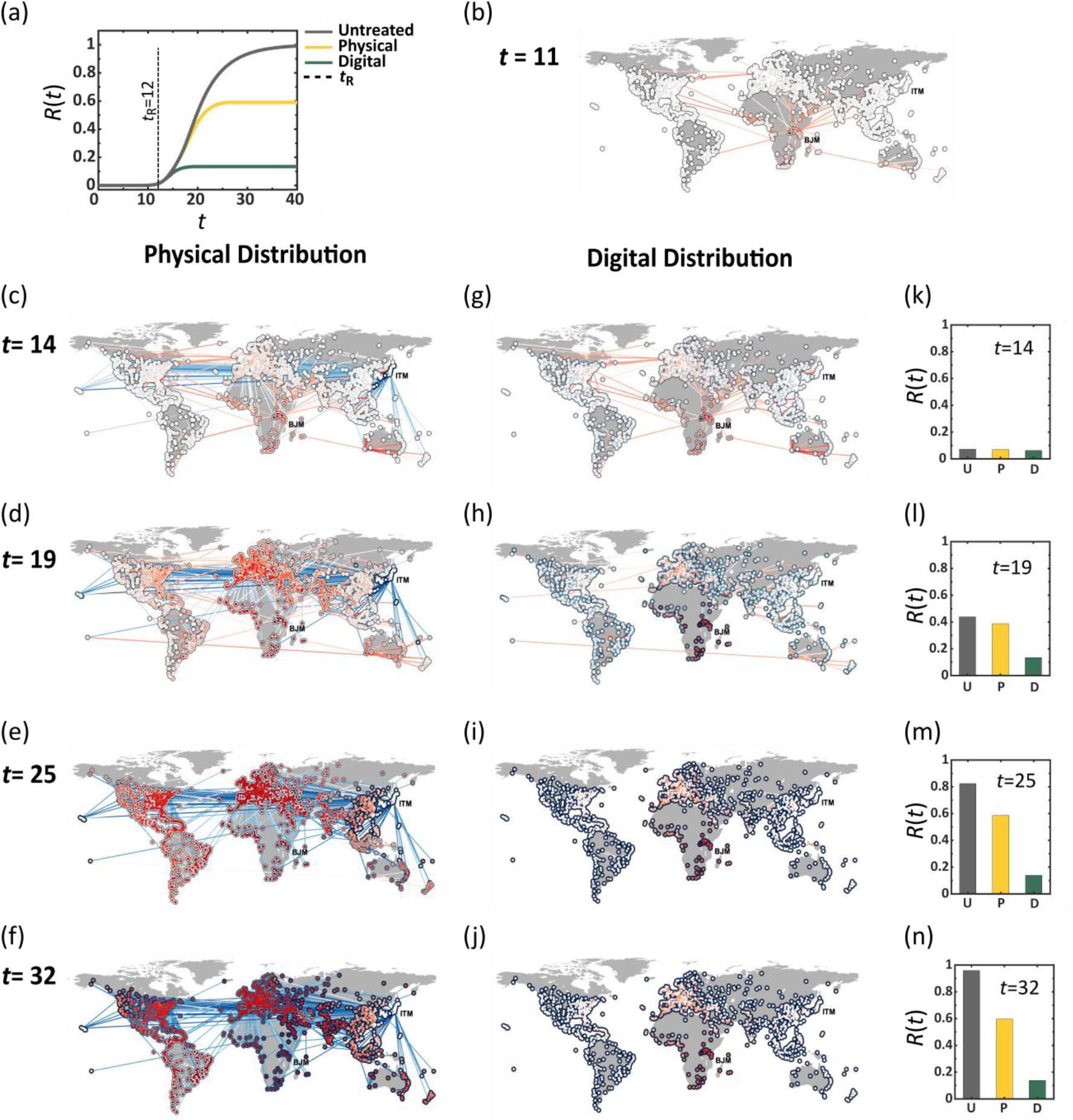
Outrunning a contageuous epidemic using Physical vs. Digital therapeutic distribution. (a) The global mortality rate *R*(*t*) vs. *t* following an outbreak at Burundi (BJM). Lacking treatment we observe *R*(*t* → ∞) ~ 1 (grey), which, following physical treatment,, initiated at *t_R_* = 12 days (dashed line), reduces to ~ 0.6 (yellow). Under identical conditions, however, digital treatment achieves a four-fold increase in effciency, reaching as low as *R*(*t* → ∞) ~ 0.15 (green). (b) The state of the epidemic at *t* = 11, directly before drug dissemination begins. The local mortality levels 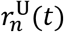 in each node and the flux of infected individuals along each link (air-route) are represented by their red color depth. (c) – (f) Physical drug dissemination begins at *t*_R_ = 12 in Osaka (ITM), with therapeutic fluxes (links) and drug availability levels (nodes circumference) represented by blue color depth. We observe a *race* between the therapeutic and the disease, both spreading along similar routes, ending in a significant fraction of deceased individuals, as indicated by the many red nodes at *t* = 32. (g) – (j) Under digital distribution each node synthesizes its own pool of therapeutics (blue circumference), resulting in a dramatic reduction in mortality, with the only *deep red* nodes, being the ones in the vicinity of BJM, that were impacted prior to our response at *t*_R_ < 12. Here we observe no therapeutic flux along the links, as the therapeutic is disseminated digitally, bypassing the physical transportation network. (k) – (n) *R*(*t*) at the four selected time points under no treatment (U, grey), physical dissemination (P, yellow) and digital dissemination (D, green). Here and throughout we set the parameters in Eq. (7) to *α* = 2 day^−1^, *β* = 0.2 day^−1^, *γ* = 1.0, *ζ* = 1.0 day^−1^, *h* = 8 and *ε* = 10^−6^. The mean capacities under both physical (10) and digital (11) distribution were set to 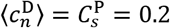 day^−1^and the response time to *t_R_*=12 days. The individual capacities 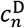 are extracted from a normal distribution 𝒩(*μ*, *σ*^2^) with mean *μ=*0.2 and standard deviation *σ*=0.1*μ.*

Following the initial outbreak at *t* = 0, we define the response time *t*_R_ as the time required to develop and begin distribution of a therapy and simulated two different therapeutic scenarios, both beginning at *t* = *t*_R_:

### Physically distributed therapy (Fig. 1c-f).

We take the classic approach, in which the developed drug is manufactured at a specific source node *s*, then distributed globally via air-transportation. In each location, some of the drug is consumed, based on the local infection levels, and the rest continues to travel to farther destinations, through pre-planned travel paths from *s* to all remaining destinations (Supplementary Section 2.2). Most crucially, the dissemination is limited by the source’s distribution capacity 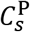 (day^−1^), capturing the number of doses that can be shipped from *s* per day, as dictated both by manufacturing capabilities and by *s*’s volume of outgoing flights. Setting 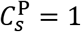 represents a scenario where *s* is capable of distributing sufficient supply to satisfy the global demand in a single day, *i.e.* produce and ship doses at a volume comparable to the entire global population. Most commonly we expect to have 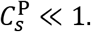

### In Fig. 1c-f

we present the evolution of the epidemic at four selected time-points. At *t* = 0 we simulate an outbreak (red) at Burundi (BJM), emulating the 2013 Ebola, which originated in Africa^33,34^, then track its spread through air-travel. The node mortality levels and the epidemic fluxes, *i.e.* the daily volume of infected passengers on each route, are represented by red color depth. Drug dissemination (blue) begins at *t*_R_ = 12 days in Osaka (ITM), again using color depth (blue) to signify the availability/flux of drugs in each node/route. The snapshots illustrate the competition between the two spreading processes – the diffusing pathogen vs. the disseminated therapeutic – showing, through the long-term prevalence of infections (red) the inefficiency of physical distribution in the face of a virulent epidemic. Indeed, the delayed response (*t*_R_), combined with the limitations of centralized manufacturing and shipping 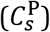 prohibit the physically distributed therapy from efficiently mitigating the pathogenic spread.

### Digitally distributed therapy (Fig. 1g-j).

Our proposed digital therapeutic is distributed as data, and hence benefits from practically immediate dissemination. Once the drug has been developed at *t* = *t*_R_, its sequence is sent to all nodes instantaneously, and synthesis is then carried out locally at each destination. Here, the main *bottleneck* for the drug dissemination is driven by the capacities 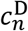 (*n* = 1, …, *N*) to locally print and distribute the digital sequence in its material form. This capacity is impacted by the abundance of printing devices in *n* and by the logistic efficiency of *n*’s health-care system in delivering the printed drugs to the infected population. Hence 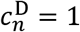 translates to a daily production and dissemination of *M_n_* doses per day, *i.e*. covering the entire local population.

For comparison purposes, note that a *mean capacity* of 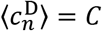 captures a state in which the distributed production covers, overall, a *C*-fraction of the global demand per day, equivalent, in terms of production capabilities to setting the physical capacity to 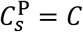. Hence setting 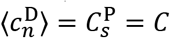 represents a scenario where both strategies, physical vs. digital, exhibit similar global production rates, thus isolating only the effect of the decentralized (digital) vs. centralized (physical) dissemination strategies.

The results of the digital therapeutic strategy are shown in **Fig. 1g-j**. As before, the spread of the disease is captured by the red nodes and links, however, in this case, the drug is no longer distributed along the same infrastructure, but rather manufactured locally at rates 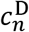, therefore, the blue links are absent. Instead, drug availability in each location is signified by the blue color depth of each node’s circumference, while infection levels are, as above, captured by the red fill of all nodes. Using similar lag *t*_R_ and capacities 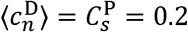, *i.e.* a daily coverage of 20%, we find that digital distribution is by far more efficient than physical distribution, in this example reducing the final mortality from *R* ≈ 1 under no treatment, to *R*= 0.15 with the digital therapeutic vs. *R* = 0.60, four times higher, under the physical therapeutic (**Fig. 1k-n**).

A crucial factor in our ability to overcome a virulent and lethal epidemic is our response time *t*_R_, required to identify the threat and begin the distribution of a therapeutic. To observe the impact of *t*_R_, we track in **Fig. 2a** the efficiency of the physical and the digital therapeutics under increasing response times. For each *t*_R_ we measured this efficiency via

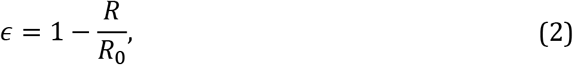

where *R* is the observed long term mortality under physical/digital treatment and *R*_0_ is the projected mortality in the absence of treatment, *i.e*. 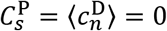. A perfect response is captured by *ϵ* → 1, *i.e. R* ≪ *R*_0_, representing a dramatic reduction in mortality, while *ϵ* → 0 indicates that mortality remained almost unchanged by our intervention. As expected we find that *ϵ* declines as *t*_R_ is increased, however for the entire range of response times the digital distribution (green) consistently achieves higher efficiency than physical (yellow). In fact, even in the ideal case, where *t*_R_ = 0, an *immediate* response, physical distribution achieves an efficiency of only 80%, while digital saves practically all potential fatalities.

**Figure 2.**
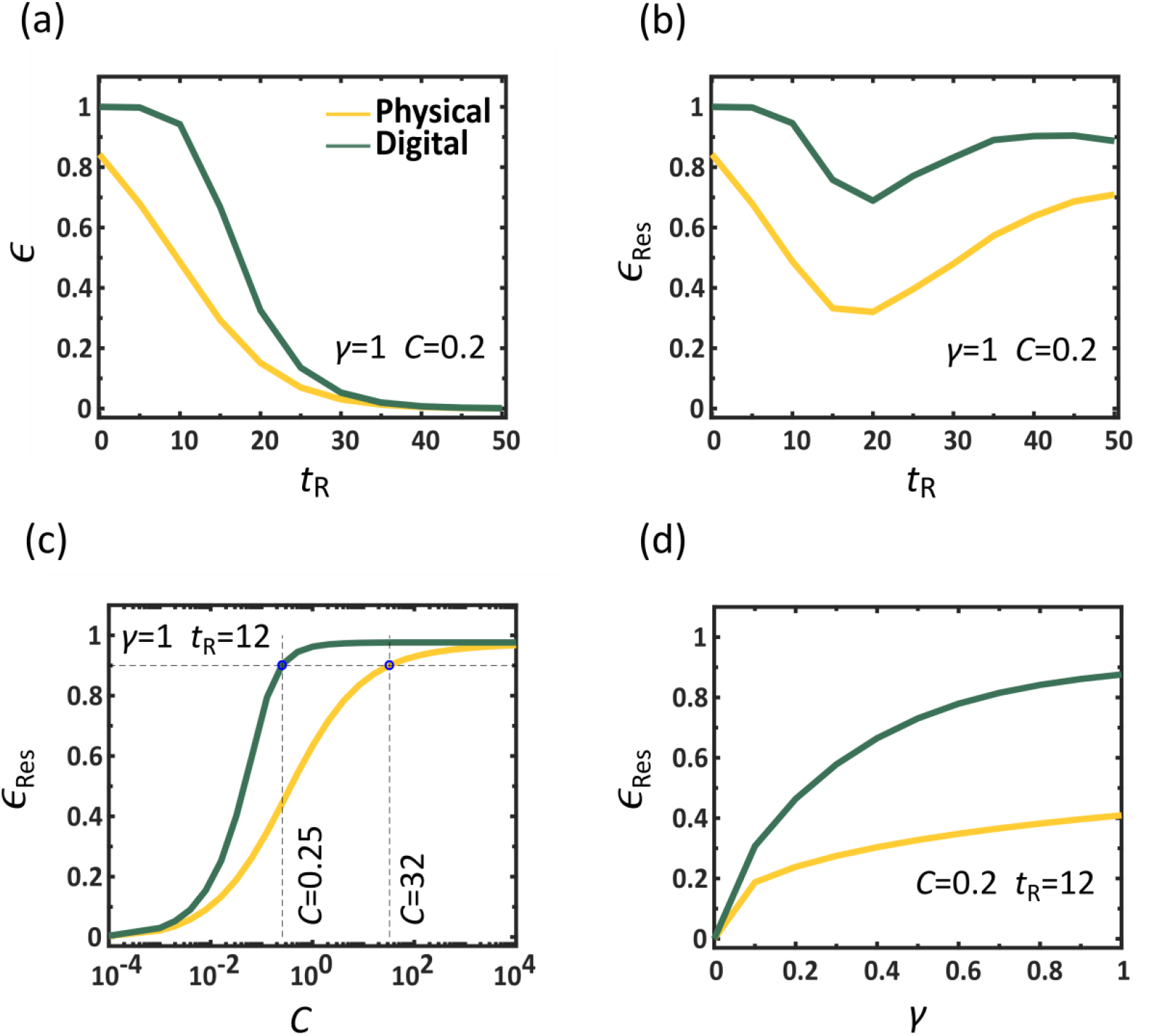
Efficiency of physical vs. digital drug dissemination. The efficiency *ϵ* vs. the response time *t*_R_ under physical (yellow) and digital (green) drug dissemination, showing that digital is consistently superior to physical. For large *t*_R_ both methods show little efficiency as the majority of the population has already been impacted. (b) The residual efficiency *ϵ*_Res_ (3) vs. *t*_R_, capturing the reduction in mortality posterior to our response, namely eliminating individuals that have already perished prior to *t*_R_. We find that even under late response (large *t*_R_), digital (green) still saves a larger fraction of the *remaining* population compared to physical (yellow). (c) *ϵ* Res vs. the capacity 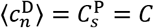 which captures the fraction of the global demand that can be manufactured and disseminated daily, either in a centralized fashion (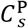, physical, yellow) or via localized production (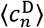, digital, green). We find that a successful mitigation under physical requires unrealistic production and shipping capacities. For instance, to achieve *ϵ*_Res_=0.9,a 90% reduction in post-response mortality, we must have a capacity of *C*>30, *i.e.* distribute doses in excess of 30 times the global demand per day (dashed lines). Strikingly, a similar efficiency can be achieved under digital therapeutics with capacity as low as *C*<0.3, representing a two orders of magnitude reduction in capacity, while achieving a comparable outcome. Therefore, digital distribution is not only faster, using data transmission instead of physical shipping routes, but also inherently, more efficient, saving more people with significantly less, and hence realistic, resources, as a result-bringing successful mitigation to within practical reach. (d) *ϵ*_Res_ vs. drug efficacy *γ.* Digital drugs achieve higher performance even if their therapeutic efficacy is low. In fact, as long as the digital therapeutic has *γ>*0.2, namely that only one out of five individuals is cured by the drug, it is guaranteed to exceed the performance of the physical treatment, even if the latter has a 100% success rate. This indicates that digital distributability, in some cases, supersedes efficacy, as a metric for ranking potential treatments. Here in each panels we varied a specific parameters(*t_R_,C* or *γ*) while controlling for all others, as appears in the panels themselves and detailed in the caption of **Fig.1.**

In the limit of large *t*_R_ both methods exhibit low efficiency, a natural consequence of the fact that the majority of the impacted population have already perished, and cannot be saved. Therefore, we consider the residual mortality Δ*R* = *R* − *R* (*t*_R_), capturing only the fatalities that occurred posterior to our response. This allows us to evaluate the residual efficiency via

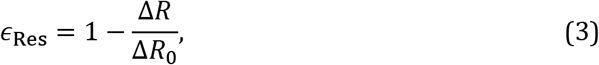

where Δ*R*_0_ = *R*_0_ – *R*_0_ (*t*_R_). We now see that even if *t*_R_ is large, our ability to save the *remaining* population is enhanced if we prioritize digital over physical distribution (**Fig. 2b**).

Next, we examine the impact of the distribution capacity on the efficiency of the disease mitigation. We consider a spectrum of capacities 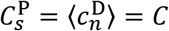, with *C* ranging from 10^−4^ to 10^4^ day^−1^, spanning a broad range, from extreme deprivation to extreme overproduction. For each value of *C* the global production and dissemination rates are identical under both digital and physical distribution, with the only distinction being the centralized production at the single source *s* (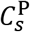, physical) vs. the decentralized synthesis in all *n* (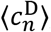, digital). We find that digital distribution is again significantly more efficient (**Fig. 2c**, green), achieving an efficiency of *ϵ*_Res_ > 0.9 already at *C*~0.3, a scenario in which the average node can only produce 30% of its demand per day (dashed-lines). Similar efficiency under physical distribution (yellow) is only achieved at *C*~30, which is not only 10^2^ times higher than the required production rate of digital, but, most importantly, an extremely unrealistic value, describing a state in which a single source node *s* produces and ships enough doses per day to cover 30 times the global demand. Optimal efficiency *ϵ*_Res_ → 1, achieved around *C* ~ 1 for digital, is only reached under the completely unattainable *C* ~ 10^4^ in the case of physical distribution.

Another crucial factor we examine is the efficacy of the therapeutic *γ*, quantifying the percentage of individuals who recover under the physical/digital treatment. Once again, we find that digital outperforms physical, achieving a higher *ɛ*_Res_, even with significantly lower efficacy *γ* (**Fig. 2d**). This demonstrates that the *efficacy* of a proposed therapeutic, the classic measure by which drugs are currently ranked^28,29^, should be balanced alongside its *distributability*, a currently ignored factor that highly prioritizes the digital strategy.

Hence, digital distribution with local synthesis is not only faster, avoiding the inevitable lag-times of physical shipping, it is also inherently more efficient, achieving, due to its decentralized nature, a dramatically higher reduction in mortality under significantly lower, and therefore realistic, production (*C*) or efficacy (*γ*) levels. Next, we show that the origins of this efficiency are rooted in the egalitarian nature of the localized production, which stands in sharp contrast to the highly unequal distribution of the physical dissemination, where the majority of doses accumulate in a small number of nodes, leaving a large fraction of the therapeutic unexploited, and a significant portion of the population deprived of the therapy.

### Distribution equality

We measured the local residual efficiency in each node as

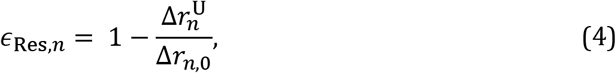

where 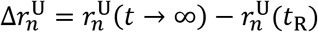 and 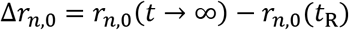 represent the residual mortality in *n* with and without the therapeutic, respectively. In analogy with Eq. (3), this local efficiency quantifies the *benefit* provided by the therapeutic to each specific location *n* on a scale ranging from zero (no benefit) to unity (optimal). This allows us to evaluate the level of benefit inequality between all nodes through the Gini coefficient^35,36^

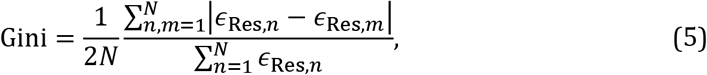

which ranges from zero, for a fully uniform *ϵ*_Res,*n*_, to unity, in the limit of extreme inequality. We find in **Fig. 3a** that under digital distribution the inequality is negligible, with Gini being close to zero independently of *C* (green). In contrast, physical distribution (yellow) creates an inherent unevenness, exhibiting a high Gini coefficient even when *C* ~1. To get deeper insight into this unevenness we calculated the probability density *P*(*ϵ*) for a randomly selected node to have *ϵ*_Res,*n*_ *ϵ*(*ϵ*), *ϵ* + d*ϵ*) for different values of *C*. In physical distribution (**Fig. 3b**) we observe for *C* = 0.01 (yellow) a high density around *ϵ* → 0, and a slight increase in *P*(*ϵ*) close to unity (see inset). This depicts a coexistence of a majority of low efficiency nodes with a selected minority of saved nodes, for which *ϵ*_Res,*n*_ is high, illustrating the severe benefit inequality. Only when *C* is set to 1 (orange) do we observe the highest density at *ϵ*_Res,*n*_ ≈ 1. Yet even under these conditions, the saved nodes continue to coexist alongside a long tail of underserved destinations whose local efficiency reaches as to low as 0.2. In contrast, under digital distribution *P* (*ϵ*) exhibits a uniform shift towards *ϵ* = 1 as *C* is increased, representing an *egalitarian* decrease in mortality, evenly spread across all populations (**Fig. 3c**).

**Figure 3.**
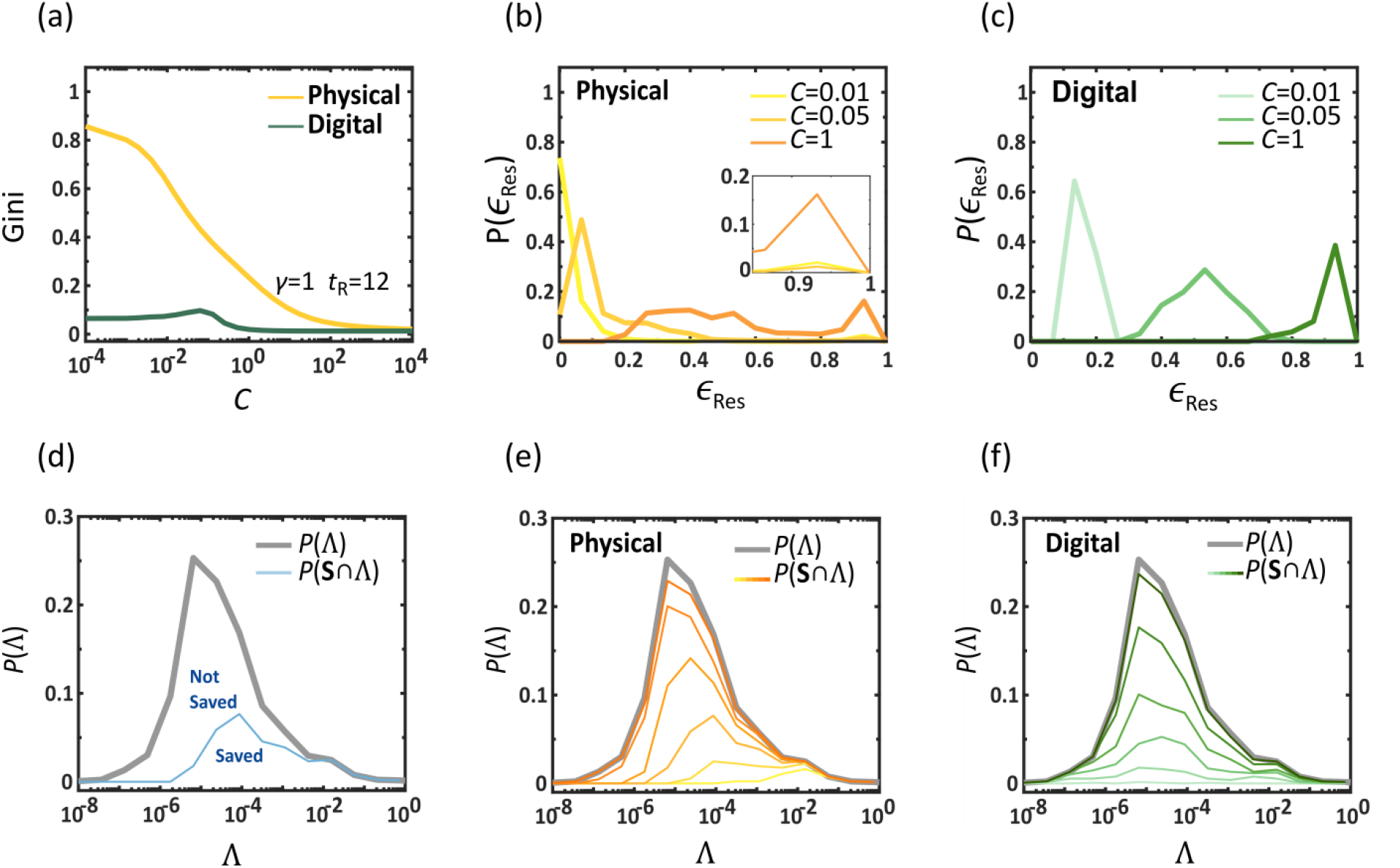
Equality and allocation of resources in drug dissemination. To understand the roots of the dramatically improved performance of digital vs. physical we examined the level of inequality in the local efficiencies *ϵ*_Res, *n*_ via the Gini coefficient (5). (a) Gini vs. *C* for physical (yellow) and digital (green) dissemination. While digital is egalitarian by nature, having a low Gini coefficient, physical is extremely uneven, with few nodes that benefit from high *ϵ*_Res_, *n* and a majority of nodes that remain deprived. This indicates that uneven allocation of resources, and not the slower shipping rates, is the main obstacle for efficient physical dissemination. (b) The probability density *P*(*ϵ*_Res_) for different capacity levels under physical distribution. We observe a coexistence of a minority of well treated nodes (peak around *ϵ*_Res_ ≈ 1, see inset) and a majority of nodes with varying efficiency levels. (c) In contrast, for digital distribution we observe a bounded *P*(*ϵ*_Res_), whose mean efficiency (peak) approaches *ϵ*_Res_ = 1 uniformly as *C* is increased. This represents an even allocation of benefits, in which most nodes witness a similar rise in efficiency as capacity is increased. (d) We measured the dilution Λ_*sn*_ in (6) and obtained the probability density *P* (Λ) for Λ_*sn*_ ∈ (Λ, Λ + dΛ). We find the Λ_*sn*_ spans several orders of magnitude, indicating that nodes receive diverse quantities of doses from *s*, exposing the roots of the uneven drug dissemination under physical distribution. Dividing the nodes into saved (**S** = 1) and unsaved (**S** = 0) we also measured *P*(**S** ∩ Λ) (blue), in which the fraction of saved nodes equals to the area under the curve. The saved nodes tend towards the lower dilution (Λ large), averting the left part of the graph. (e) *P*(**S** ∩ Λ) as obtained for increasing capacity *C* (yellow to orange, C = 0.01, 0.1, 0.3,1.0, 3.0, 10 days^−1^). While the number of saved nodes (area) increases with *C*, it features a systematic bias towards low dilution, an intrinsic inequality rooted in the structure of the distribution network *B_nm_*. (f) Under identical conditions, digital distribution also shows a gradual increase in saved nodes (area under *P*(**S**∩ Λ), shades of green), however, in the absence of a distribution *network*, the saved nodes are evenly spread, independently of Λ (light to dark green, *C* = 0.025, 0.03, 0.035, 0.04, 0.05, 0.1 days^−1^).

To understand the origins of this unevenness consider the routing of the physical therapeutic through the shipping fluxes *B_nm_* and *Z_nm_* in Eq. (10). Unless we intervene and take control over the air-transportation network, an unlikely and economically harmful scenario, these fluxes reflect the *natural* flow of passengers and cargo through the existing air-routes^18,23^. Under these conditions, for every *q_m_*(*t*) doses present in *m*, a fraction *B_nm_* will be shipped throughout the day to *n*, then yet a smaller fraction will propagate onwards to *m’s* next neighbors and so on. Hence the therapeutic availability becomes diluted as if flows downstream from the source *s* to the target *n*. Accounting for all pathways from *s*, the *dilution* at *n* becomes (Supplementary Section 2.3)

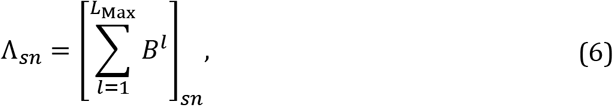

namely the *s,n* term of a geometric series with base *B*. Equation (6) captures the therapeutic dilution as it flows downstream from *s* to all target nodes *n*. Roughly speaking, it approximates the number of doses reaching *n* per each dose exiting *s*, hence indicating which nodes *n* benefit from superfluous drug availability (large Λ) and which will be underserved (small Λ).

To observe the impact of this diversity we constructed *B_nm_* from the empirical fluxes of human mobility (Supplementary Section 2.2) and used (6) to obtain the dilution Λ_*sn*_ of all nodes *n*. In **Fig. 3f** we show the probability density *P*(Λ) for a randomly selected node to have Λ_*sn*_ ∈ (Λ, Λ + dΛ); grey solid line. We find that the dilution spans orders of magnitude, indicating an extremely uneven distribution: for each unit of therapeutic leaving *s*, a small minority of nodes, whose Λ_*sn*_ is of order 0.1 − 1, receives a major fraction, whereas an overwhelming majority of nodes, whose Λ_*sn*_ ~ 10^−5^ − 10^−7^, remain with only a tiny fraction of the dosage originating in *s*. Therefore, to reserve sufficient quantities of doses, *i.e. q_n_*(*t*) ~1, for these deprived nodes, which, by far, represent the majority of potential destinations, *s* must have an inconceivable capacity of 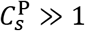, as indeed was found in **Fig. 2c**.

This diversity in Λ_*sn*_ directly impacts the probability of nodes to be *saved* by the distributed therapeutic. For a node *n* to be considered saved, *i.e.* **S**_*n*_ = 1, we require a significant reduction in its mortality, namely that *R_n_*/ *R_n_*,_0_ < *η*, where *R_n_* and *R_n_*,_0_ are the long term mortality rates in *n* with and without treatment, respectively. Setting *η* = 0.5, we measured *P*(**S** ∩ Λ), the probability that a randomly selected node in the group Λ_*sn*_ ∈ (Λ, Λ + dΛ) is saved. As expected, we find that the least diluted nodes, *i.e*. ones with large Λ_*sn*_, have the highest probability to be saved (**Fig. 3e**, shades of orange). As 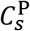 is increased, the fraction of saved nodes, namely the area under *P*(**S** ∩ Λ) also increases, but the preference towards large Λ_*sn*_ continues to underlie the uneven treatment pattern, with **S** consistently concentrated around the least diluted nodes. Repeating this examination for digital distribution (**Fig. 3f**) shows the intrinsic difference between the two strategies, as here, since *B_nm_* plays no role in delivering the therapeutic, *P*(**S**∩ Λ) rises uniformly across all nodes, independently of their highly uneven pathways to the source *s*.

Our findings expose an intrinsic lacuna in the air transportation network, that, by design, leads to an extremely unequal distribution, and hence to a highly inefficient and discriminative spread of the therapeutic. We have shown this here on *B_nm_* constructed from the *natural* fluxes, as extracted from global aviation data, assuming little intervention in flight routes and shipping capacity. One may, indeed, consider more efficient distribution networks, as we do in Supplementary Section 2.2, however, under any reasonable construction, dilution in the downstream flow of the therapeutic in the form of (6) is practically unavoidable. Especially if the disease spreads globally, and we must stretch all available aviation resources to ship the therapeutic worldwide – conditions rendering dilution inevitable.

In Supplementary Section 2.2 we also construct an *egalitarian network*, which indeed rectifies, to some extent, the observed distribution inequality. However, the realization of such a network requires us to seize full control over air transportation, which is not only an unrealistic cenario, but also, as mentioned above, one that imposes an extremely heavy burden on the global economy and political stability.

### Optimizing physical distribution

Our modeling up to this point was based on a diffusive propagation of the physical therapeutic, *a la* Eq. (10), a framework that can be naturally coupled to the SIR epidemic spread of Eq. (7). In this framework our control over the distribution efficiency is enacted through the design of the networks *B_nm_* and *Z_nm_* (Supplementary Section 2.2), however, once these are set, the spread of the therapeutic is governed by diffusion, which is often sub-optimal, allowing, *e.g.*, for superfluous quantities to accumulate at selected locations. We can improve distribution efficiency by modeling it as a commodity flow problem^37,38^, seeking to optimally utilize the routes of the existing air-traffic network, until meeting the demands of all destinations^39,40^. In this framework, each air-route is assigned a carrying capacity *C_nm_*, capturing the number of doses it *can* transport per day, and each destination *n* is assigned an initial demand *d_n_*(0), depending on the size of its local population. Each step (day), as the therapeutic is shipped and accumulates at *n*, the local demands are updated, *d_n_*(*t*), subtracting the supplied doses from *d_n_*(0), until their quota is filled at time *tn*, *i.e.* the time when the remaining demand in *n* is *d_n_* (*t_n_*) = 0 (Fig. 4a). Using linear optimization, we derive the optimal dissemination strategy to achieve maximum daily flow to all destinations *n* = 1, …, *N* from the source *s*, avoiding any wasted dosage via oversupply, and satisfying the constrained carrying capacities *C_nm_* (Supplementary Section 4).

**Figure 4.**
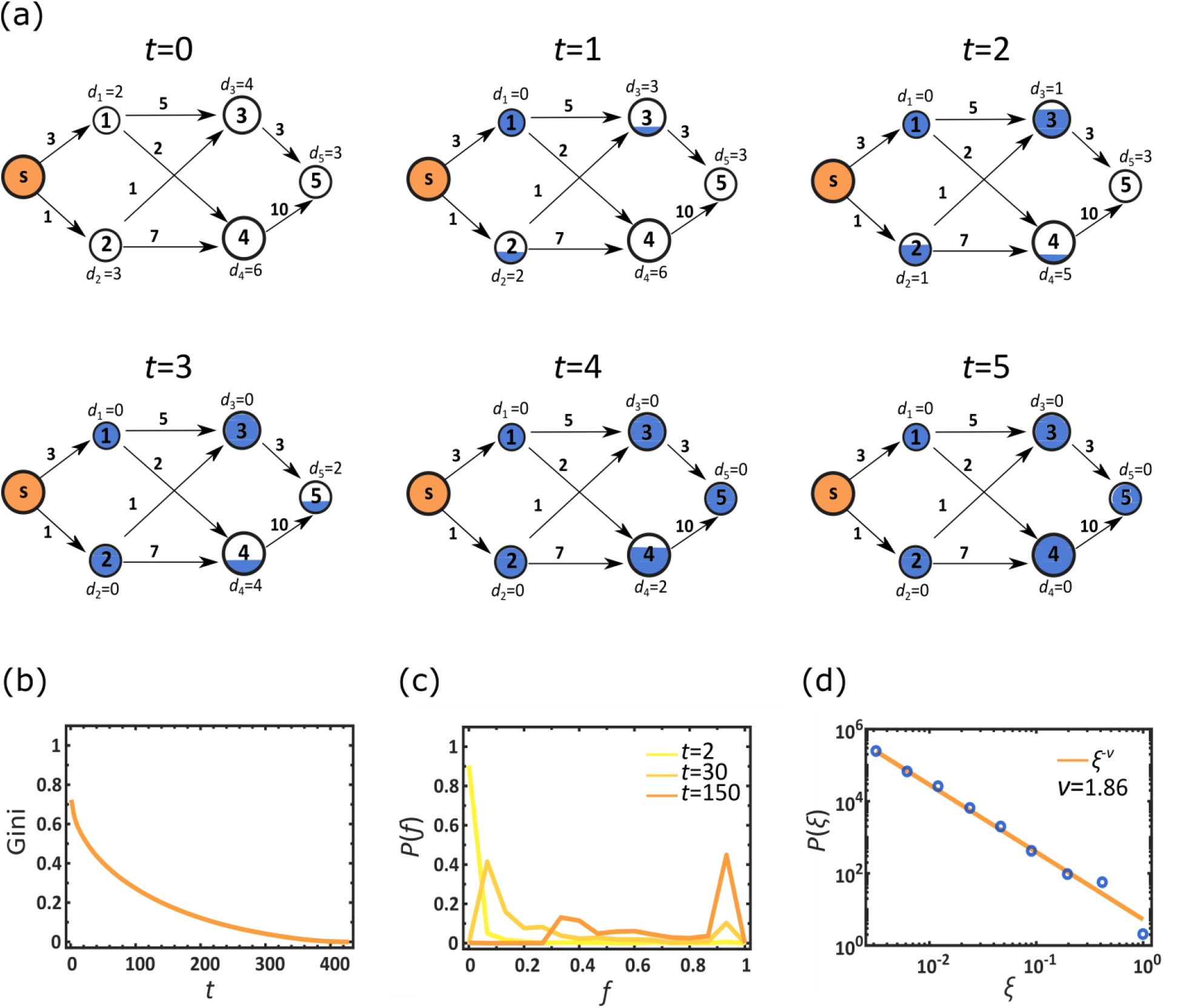
Optimizing physical distribution. (a) To enhance the efficiency of physical distribution we used the commodity flow framework: each node is assigned an initial demand *d_n_*(0), proportional to its population (node size); and each air-route is assigned a daily carrying capacity *C_nm_* (numbers by edges), proportional to its daily volume of passenger traffic. In each time step *t* we optimize the flow from the source *s* to supply as much of the therapeutic as possible, within the constrained *C_nm_*. As nodes accumulate the therapeutic (blue fill) their demand is updated accordingly. For example, node 2’s initial demand is *d*_2_(0) = 3 units. Following the first round of shipment (*t* = 1) a single unit is shipped from *s*, supplying a fraction *f*_2_(1) = 1/3 of *d*_2_ (0), and hence reducing its demand to *d*_2_ (1) = 2. A node is fully supplied (fully blue) at time *t_n_*, when *f_n_*(*t_n_*)= 1, and accordingly *d_n_*(*t_n_*)= 0. For node 2 we have *t*_2_ = 3. (b) The Gini coefficient vs.*t*, as obtained from the time dependent fractional stock levels *f_n_*(*t*). We find the despite the optimization, inequality continues to govern the therapeutic distribution, as expressed by the high Gini coefficient at small *t*. This describes a scenario in which few nodes are fully stocked early on, while the majority of nodes will only reach *f_n_*(*t*) → 1 (and hence Gini → 0) at much later times. (c) The probability density *P*(*f*) vs. *f* at three selected time points *t*. Similarly to Fig. 3b we observe a highly unequal distribution in which selected nodes are fully supplied (right peaks), while others are still deprived (left peaks). These patterns in (b) and (c) are strikingly reminiscent of the those observed earlier in Fig. 3a,b. (d) *P*(*ξ*) vs. *ξ*, capturing the probability density that a randomly selected node enjoys a supply rate of *ξ* ∈ (*ξ* + d*ξ*). The power-law form of *P*(*ξ*) (solid line represents *ξ* ^−*v*^) demonstrates the uneven dissemination of the therapeutic, in which supply rates range over orders of magnitude.

Our previous analysis in **Fig. 3** indicated that the root problem in physical dissemination is its extreme levels of inequality, as expressed through the efficiency *ϵ*_Res, *n*_ in Eq. (4). In the context of the current modeling this is most naturally expressed through

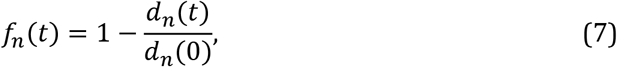

capturing the fraction of *n*’s demand supplied by the time *t*. Interestingly, we find in **Fig. 4** that the optimal commodity flow, in spite of being profoundly different from the diffusive propagation of Eq. (10), leads to strikingly similar patterns of unevenness. For instance, the Gini coefficient extracted from *f_n_*(*t*) remains large at the early stages of the dissemination (**Fig. 4b**), reminiscent of the patterns observed for *ϵ*_Res,*n*_ in **Fig. 3a** above. This indicates that few nodes fill their initial demand early on, while the majority of nodes take a long time to satisfy their quota, hence the large inequality observed for small *t*. Similar patterns are also observed by the time evolution of *P*(*f*), capturing the probability density for a random node *n* to have *f_n_*(*t*) ∈ (*f*, *f* + d*f*); **Fig. 4c**, as compared to **Fig. 3b**. Indeed, *P*(*f*)recovers the signature two peak structure observed for *P*(*ϵ*_Res_), with increased density around *f* → 0 and *f* ~ 1, a coexistence of early vs. late supplied nodes. Finally, we measured the supply times *t_n_* for all nodes *n* to fill their demand, extracting each node’s *supply rate *ξ*_n_* = 1/*t_n_*, *i.e.* the average volume of doses entering *n* per unit time. In **Fig. 4d** we find that *ξ_n_* follows a power law distribution with *P*(*ξ*) ~ *ξ*^−v^ (*v* ≈ 1.86), precisely capturing the predicted inequality, in which a majority of underserved nodes (small *ξ*) coexist with a small minority of nodes whose supply rate is orders of magnitude above average.

Together we find that even under optimal distribution, the unequal supply rate, indeed the root cause of the inefficiency of the physical therapeutic, is practically impossible to avoid. Therefore, the lacuna of physical distribution is not unique to our modeling via Eq. (10), but rather represents an intrinsic property of the existing transportation networks, thus illustrating the crucial need for a decentralized strategy for drug dissemination.

## DISCUSSION

Our analysis outlines a currently unexplored strategy to address the threat of global epidemics. It is not only more efficient than the standard response protocol, but, in the face of a lethal and highly contagious epidemic, it is likely the only viable option. We considered two potential realizations of this strategy, based on nucleic acids or peptides, which, in our view, represent the most promising candidates for digital therapeutics, given the current technology (**Box I**). Other digitizable media, if become available, are equally relevant. Indeed, our purpose here is to expose the digital mitigation *strategy*, rather than to promote specific types of therapeutics. Still, of the currently available candidates, and specifically between nucleic acids vs. peptides, we believe that DNA aptamers exhibit several advantages^41^, such as the diversity of their initial random library^42^, their rapid discovery process^43^, and their relatively stable nature, which underlies their smooth handling and shipping, compared to peptides.

Aiming towards application, however, we must first address several technological gaps pertaining to the scalability of DNA printing^41^. Present day oligonucleotide synthesizers are capable of synthesizing ~ 1 gram/day of a short oligonucleotide, at a cost of ~ 50 USD, requiring thorough purification, and generating, as a byproduct, large amounts of liquid waste. Combined with the current cost of the required instrumentation, estimated at ~ 250K USD, we believe, that despite the unequivocal advantages of digital distribution, it is currently of limited applicability at a global scale. This status-quo, however, is a consequence of our current priorities, rather than an intrinsic technological restriction. Indeed, the need for scalable low-cost mass-printing of short DNA sequences was not evident until now, and hence the development of the relevant technology was never prioritized. Our findings here suggest reassessing these priorities^44^.

Another crucial issue regarding the mass-production of DNA-based therapeutics, is the need for large amounts of phosphorous, a limited resource, that constitutes ~ 10% of the DNA mass^45^. It was recently estimated that Earth’s accessible phosphorous reserves are in excess of 6 × 10^13^ kg, with global production in 2016 amounting to 2.6 × 10^11^kg, mainly serving the fertilizer industry^46^. Our calculations, based on these figures, indicate that phosphorous availability is orders of magnitude higher than that required for global dissemination of our proposed DNA-based therapeutics. To be on the safe side, we assume the oligonucleotide dosage to be of the order of ~ 10^−2^ kg per person, and the synthesis yield to be only 10%. Even under these stringent assumptions, we can treat the global population of ~ 10^10^ individuals with ~ 10^8^ kg of phosphorous, amounting to less than 0.1% of the current annual consumption.

We tested both approaches – digital and physical - under similar terms: a rapidly spreading epidemic, a similar response time *t*_R_, similar drug efficacy *γ* and comparable capacities 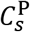 and 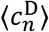, therefore singling out the specific impact of the distribution method, finding the consistent superiority of digital over physical therapeutics. In reality, however, digital distribution benefits from several additional advantages. Consider, for example, 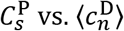. The former captures the capacity of a specific node *s* to manufacture and spread a therapeutic in quantities that can meet the global demand. In a worldwide pandemic this scales with the global population, likely exceeding the capacity of any one node. In contrast, under digital distribution each node synthesizes the therapeutic locally, having only to produce enough for its own population’s demand. If we indeed develop the technology to mass-print short DNA strands, it is likely that highly populated nodes, will also have an increased capacity to synthesize the drugs, *e.g.*, by scaling the number of printers with population size. Therefore, while typically 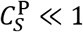, confronting a single node vs. the global demand, 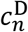 is likely of order unity for most *n*. Moreover, in real scenarios, nodes with 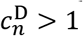 can aid less capable nodes by small scale per demand physical distribution – a practical strategy to further improve dissemination efficiency, that was not included in our current theoretical framework. Finally, in case of a novel pathogen, the discovery of a therapeutic aptamer is often significantly faster^43^ than that of molecular drugs (**Box I**), potentially reducing our response time *t*_R_, which as our analysis indicates, plays a crucial role in the mitigation efficiency.

### Box I. Digitizable therapeutics

While classic medications are shipped and transported in their physical form, a *digitizable* therapeutic can be distributed as data and manufactured on location via scalable printing technology. Such distribution can be potentially achieved using polymer-based therapeutics, natural or artificial, taking advantage of their discrete recurring building blocks, which can be naturally mapped into a data file detailing the polymer sequence. Two most relevant examples of such polymers, potentially employed as digital therapeutics are nucleic acids and peptides, whose therapeutic potential is already demonstrated on several existing treatments, such as Fomivirsen^47^, Pegaptanib^48^ and Insulin^49^.

#### Nucleic acids

Built from a selection of four natural building blocks (G, C, A, T/U) and additional artificial ones, DNA/RNA sequences operate either by interfering with gene expression, *e.g.*, antisense oligonucleotides^50–52^ and RNAi^53–55^, or through their unique folding geometry, which allows them to interact structurally with a target molecule, *e.g.*, aptamers^41, 56–60^ and ribozymes^61–63^. While not widespread at this point, these short DNA-sequences have already proven their potential efficacy, for example inhibiting the activity of human immunodeficiency virus (HIV)^64,65^, as well as other applications at various stages of clinical trials^58^. A crucial advantage of DNA aptamers is their rapid *ab-initio* discovery, which can be accomplished within hours or days^43^, allowing a potentially rapid response when confronted with an unknown pathogen. Their synthesis is sufficiently facile and inexpensive as to be carried out locally^41,66^, and they typically require no further chemical modification to become functional, strengthening their potential utility as digital therapeutics.

#### Peptides

Peptide sequences are coded from a selection of twenty natural building blocks, *i.e.* amino acids, folding into a defined geometric 3D structure to bind or modify selected target molecules^67–69^. Short peptides can be efficiently digitized, thanks to their relatively facile printability^70^; larger ones, however, often require additional modification (*e.g.*, S-S bonds)^71^, and are hence harder to produce and fold properly. Finally, proteins, essentially very large peptides, are at present too complex for local synthesis outside a biological system, such as a cell culture, therefore prohibiting their current use as digital therapeutics.

### Box II. Modeling a lethal global epidemic under physical and digital therapy

In a network of *N* coupled nodes *n* = 1, …, *N*, each with a population of *M_n_* individuals, we use the SIR model to track the fraction of *M_n_* who are susceptible (*s_n_*), infected (*j_n_*) or removed (*r_n_*). Each of these sub-populations is divided among the treated individuals 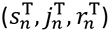, who have been provided a therapeutic, and the untreated individuals 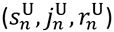, who have not yet gained access to it. For a lethal epidemic, 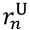 represents the deceased population, while 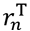 are the *saved* individuals, who, absent any treatment, would have perished. The epidemic dynamics is driven by (**Supp. Sec. 1**)

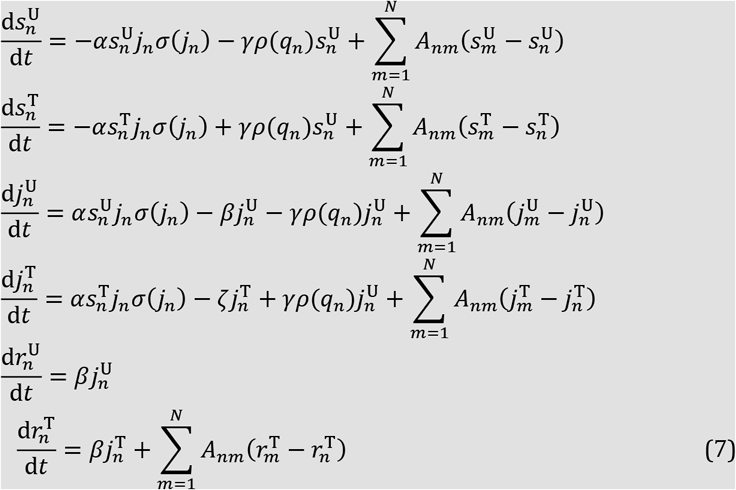

where *α* is the infection rate, *β* is the mortality rate of the untreated individuals and *ζ* > *β* is the recovery rate under treatment. The therapeutic efficacy is 0 ≤ *γ* ≤ 1 and its consumption rate *ρ*(*q_n_*) depends on the availability of the therapeutic *q_n_*(*t*) in *n*, as

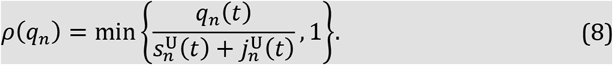

Hence *ρ*(*q_n_*) increases linearly with *q_n_*(*t*) as long as the demand (denominator) exceeds the supply, and saturates to unity when *n* has excess quantities of the therapeutic, avoiding over consumption (**Supp. Sec. 1.2**). In (7) we introduced an invasion threshold *ɛ* through the sigmoidal function

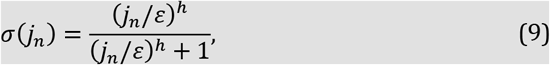

which activates the local SIR dynamics only when the local infection levels 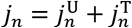 exceed *ε*. The diffusion of individuals between nodes is mediated by *A_nm_*, derived from the empirical international air-travel network^61^ (**Supp. Sec. 2.1**).

#### Drug availability

Under *Physical distribution* we have

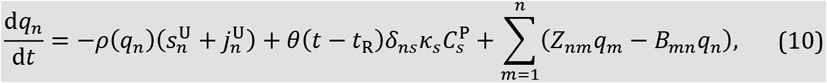

where the first term captures local drug consumption, and the second term, activated following a response time *t*_R_, represents drug production/shipment from the source *s*; *δ_ns_* is the Kronecker *δ*-function and *θ*(*x*) is the Heavyside step-function. The pre-factor *κ_s_* is derived in **Supp. Sec. 1.3**. The shipping routes are governed by *Z_nm_* and *B_mn_*, constructed in **Supp. Sec. 2.2**.

Below and in **Supp. Sec. 4** we also consider optimal physical distribution strategies based on commodity flow optimization.

In *Digital distribution* we have

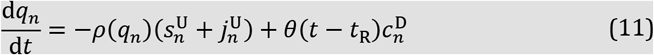

in which the central production in *s* is replaced by local production in each node at a rate 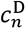. Here 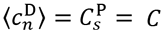 translates to a cumulative production capacity of a *C*-fraction of the global demand per day, with the only distinction being whether this production is centralized 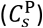 or distributed 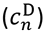; **Supp. Sec. 1.3**.

